# Feminization of the precarious at the UNAM: examining obstacles to gender equality

**DOI:** 10.1101/2024.05.13.593992

**Authors:** lu ciccia, Geraldine Espinosa-Lugo, Graciela García-Guzmán, Jaime Gasca-Pineda, Patricia Velez, Laura Espinosa-Asuar

**Affiliations:** Centro de Investigaciones y Estudios de Género, Universidad Nacional Autónoma de México, México; Facultad de Filosofía y Letras, Universidad Nacional Autónoma de México, México City, México; Secretaría Técnica, Instituto de Ecología, Universidad Nacional Autónoma de México, México City, México; Departamento de Ecología Evolutiva, Instituto de Ecología, Universidad Nacional Autónoma de México, México City, México; Departamento de Botánica, Instituto de Biología, Universidad Nacional Autónoma de México, México City, México

**Keywords:** gender gap, STEM, labor precarization, vertical segregation, horizontal segregation

## Abstract

The STEM workforce is marked by the persistent underrepresentation of women. Herein we seek a better understanding of this gender gap in different science disciplines within Latin America. Specifically, we analyzed the professional development of women in science research institutions of the National Autonomous University of Mexico (UNAM). Our analysis deepened how horizontal and vertical segregation are combined with symbolic and structural obstacles, as well as economic labor precariousness, within the framework of gender norms. We explored shared trends in the Global North to explore the perpetuation of gender stereotypes in the production of scientific knowledge; and investigated the relationship between the values implied in gender norms and the cultural capital represented in each of the disciplines. As a result, we evidenced that the knowledge areas that currently represent the highest cultural capital (pSTEM) mainly pose symbolic (and less structural) obstacles for women. Conversely, women in fields with less masculine-coded values face mainly structural barriers. This information sheds light on the gender biases that exclude women from STEM.

## Introduction

### STEM and pSTEM disciplines: analytical context

The acronym STEM (Science, Technology, Engineering, and Mathematics) was introduced to the United States by the National Science Foundation in the year 2000 to encourage interest in the disciplines it encompasses [1]. This promotion arises from the increasing demand for scientific and technological education, driven by the constant changes in technology and the growing digitalization of contemporary societies [2]. Nonetheless, workforce representation in STEM is characterized by a strong gender disparity, with the under-representation of women [3].

Systematic research addressing the underrepresentation of women in science and technology predates the formalization of the STEM acronym [3]. In recent years, there has been a marked increase in studies examining the challenges and barriers faced by women in science; for instance, in the last decade around 15,000 articles covering various disciplines such as psychology, economics, biology, physics, education, mathematics, and more have been published [4].

In the context of such disciplinary diversity, the gender gap within STEM has become especially notable due to the ‘horizontal segregation’ that characterizes these disciplines. Given the high demand for STEM workers [5] and the focus on “digitally driven economies” [2], socioeconomic status intersects with the STEM gender gap. Herein we follow the distinction between STEM and pSTEM disciplines to specify a subset of STEM fields, primarily focusing on physics [6], and to address the gender gap in physics-related disciplines, distinguishing it from other STEM areas where women might have more equitable representation. This approach acknowledges that the challenges and barriers women face may differ across various STEM fields.

Published literature on women in pSTEM is still insufficient to attain a comprehensive analysis of gender bias in science disciplines. Specifically, the relationship between women and STEM in Latin America has been neglected, with some exceptions [7]; and comparative analyses aiming to understand differences between STEM departments in Latin American universities remain to be elucidated. This approach is essential for two reasons: 1) to contribute to a better understanding of the gender gap regarding the reality of women in the different disciplines contained in STEM in Latin America, and 2) to analyze commonalities regarding the situation in the Global North. These points are essential in understanding how gender norms operate to exclude women in specialized academic contexts.

Our research efforts focused on the National Autonomous University of Mexico (UNAM). Previous works at this university [8] have demonstrated gender disparity regardless of the STEM disciplinary area. In addition, the academic careers of women in this university face significant obstacles in contrast to men who achieve seniority in a shorter period, meaning that female researchers are less represented at the highest levels of appointment and in the field of scientific research in general. The combination of both facts results in a marked segregation of women [8].

### An exploration of the obstacles underpinning gender equality

This study aims to untangle this gender gap in three research institutes of the UNAM that belong to the fields of Exact and Natural Sciences. We conducted a comparative analysis that considers how horizontal segregation (fewer women in occupational areas considered ‘masculine’) and ‘vertical segregation’ (the absence of women in senior posts) interact, and, at the same time, their relationship with the socio-economic status of academic personnel. To do this, we selected a representative pSTEM institution, the Institute of Mathematics, and two other institutions of STEM, where, due to the increasing number of women, less horizontal segregation is observed: the Institute of Biology and the Institute of Ecology (we will refer to those other areas that are not pSTEM simply as STEM, understanding that this is a broader category that we will use to encompass some Natural Sciences unrelated to physics and similar areas). Additional data from five pSTEM and six other STEM UNAM institutes was analyzed to corroborate trends observed within the three selected institutes.

Traditional views assume that women face numerous disadvantages in institutions where they are underrepresented. In contrast, we postulate that although there are few women in the pSTEM, they are less affected by vertical segregation than STEM women. This hypothesis is grounded in the observation that pSTEM fields are often seen as embodying values traditionally associated with masculinity, such as objectivity, neutrality, abstraction, reason, and universality [9–11], having the highest economic and cultural capital. So, values usually associated with femininity, such as emotion, sensitivity, and empathy, are typically excluded [12–14]. Consequently, they incorporate symbolic barriers that hinder women from entering these spaces. We propose that once these barriers are partially subdued, women’s progress is comparable to that of men entailing fewer complications. In other words, due to the symbolic barriers that discourage women from engaging with pSTEM disciplines, horizontal segregation could mitigate vertical segregation.

We hypothesize that the gendered gap will differ across scientific disciplines, depending on the values each one embodies, and the consequent economic and cultural capital associated with them. In our view, horizontal and vertical segregation intertwine with the symbolic and structural barriers that women face in our androcentric societies. We evaluated in detail the relationship between segregation and the values associated to the pSTEM and STEM disciplines. This was analyzed in terms of their being disciplines more or less masculinized and, consequently, their level of cultural capital (i.e., the epistemic authority implied in the knowledge generated in each discipline, including their social and economic status), and the level of precarization among female academic staff. Our analyses revealed that reducing horizontal segregation in fields like ecology and biology can exacerbate vertical segregation. This occurs because, while these fields may be less masculinized (valuing direct interest in life and the environment), the resulting knowledge often carries less economic and cultural capital. Consequently, gender inequality persists as men tend to occupy decision-making positions, --entailing social status and economic income--[4, 15, 16], reinforcing the idea that epistemic authority in these fields is tied to the academic status of researchers. Specifically, reduced horizontal segregation can reveal clearer structural obstacles (concrete barriers stemming from gender norms and stereotypes) that hinder the advancement of female staff to senior appointments, even when equally qualified.

## Results

The total academic population of the Institute of Biology (IB) has 71 faculty members with feminine names (f) and 90 with masculine names (m). Particularly, we registered 75 faculty members (24f, 51m) as research personnel. For the Institute of Ecology (IE), we recorded 44 f and 36 m, out of which 45 (23f and 22m) corresponded to the category of research personnel. Lastly, in the Institute of Mathematics (IM), we calculated 28 f and 88 m, out of which 95 (22f and 73m) were identified as research personnel (S1 Table).

The average age of research personnel for the three institutes is similar (IB = 56.57, IE = 56.81, IM = 56.65). The two institutes with the largest difference in the average age between feminine and masculine are IE and IM (6 - 7 years apart); IB presents a difference of approx. 2 years (S1 Fig). It is worth noting that feminine or masculine age distribution did not present obvious differences for the three institutes (S2 Fig). In terms of economic remuneration, the highest level for technical personnel is equivalent to the lowest position for research personnel (S2 Table).

Concerning the proportional distribution of the highest levels of Academic and Incentive Programs (A&IP) for the research personnel (Fig 1, S3 Fig) we observed that the IE has the greatest gender differences for the “Titular C” academic position and for the highest position in the incentive programs (PRIDE D and SNI III), whereas the IM showed an equivalent proportional distribution, but lower in SNI III level. Regarding the Emeritus honorary title (within both academic positions and the SNI incentive program), we did not register feminine names, except for the SNI in IM (Fig 2).

**Fig 1.**
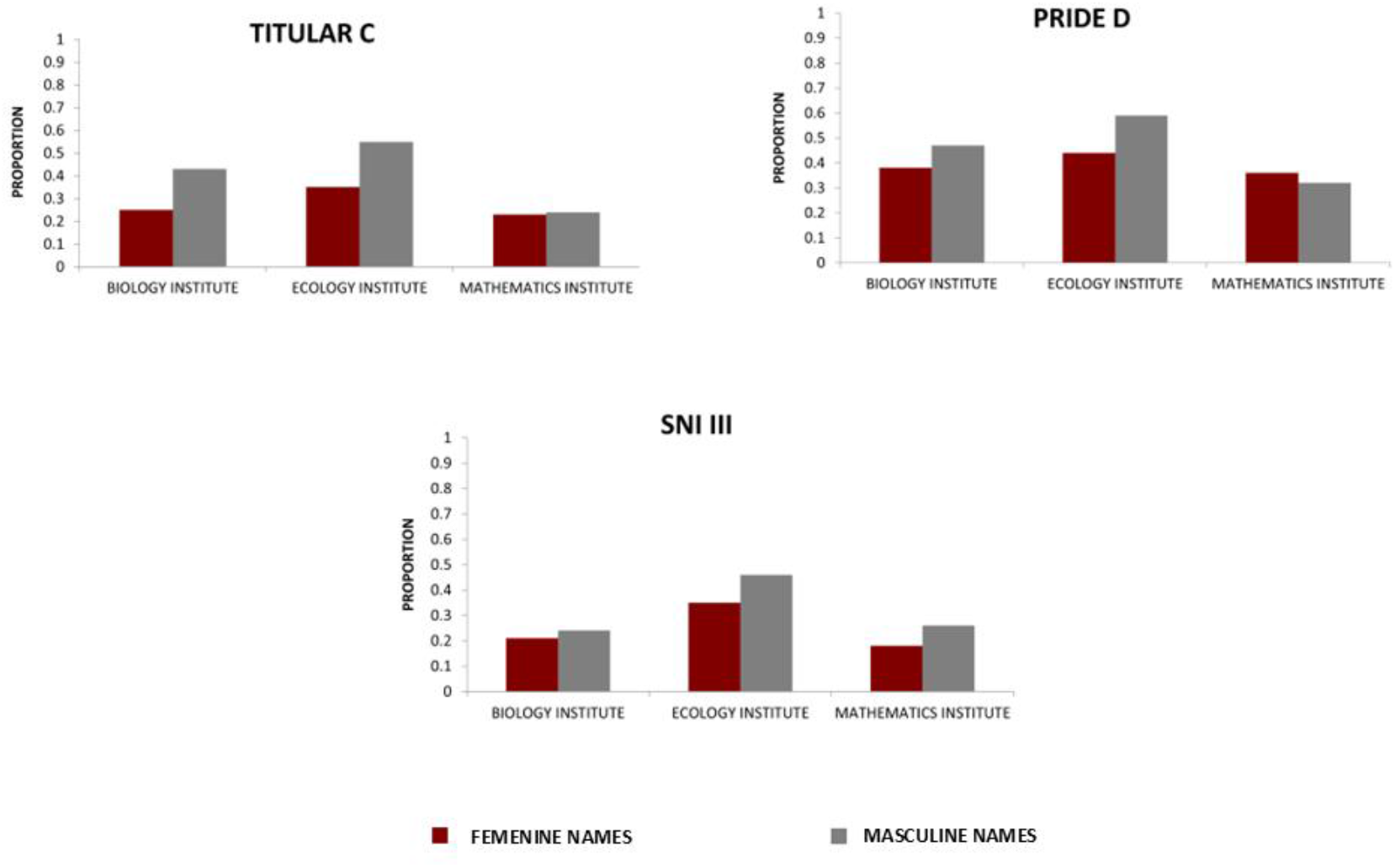
Proportional contribution comparison in the three UNAM institutes. Comparison was made for the highest levels A&IP positions (feminine vs masculine research personnel), for Biology, Ecology, and Mathematics institutes. The whole proportional contribution for all levels is shown in S3 Fig.

**Fig 2.**
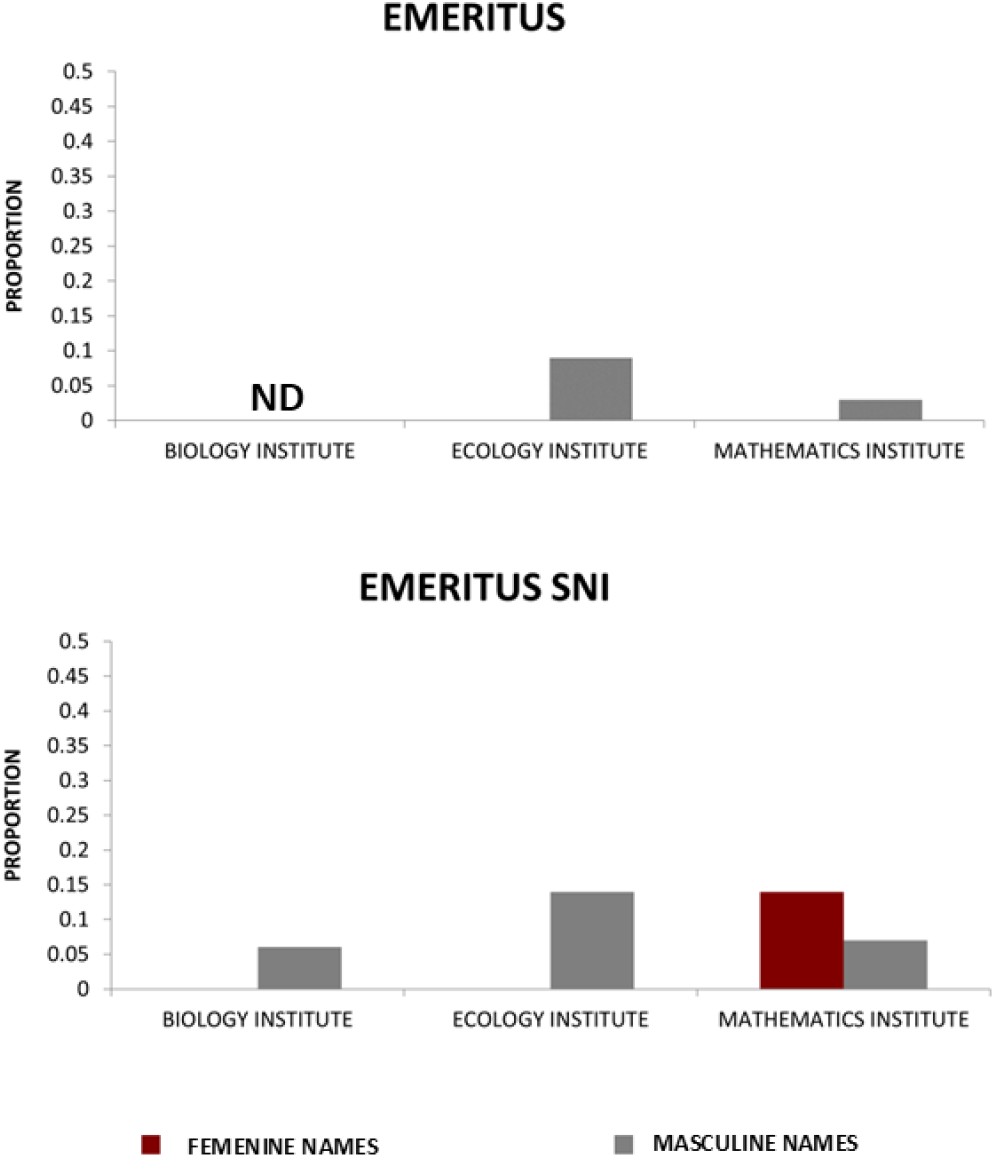
Proportional contribution of feminine and masculine research personnel occupying Emeritus distinction. Both academic positions and SNI incentives within the three institutes are shown. The whole proportional contribution for all levels is shown in S3 Fig.

Based on the gender distribution of further analyzed institutes (five pSTEM and six STEM, see details in Methods) we identified both patterns: marginally feminized and masculinized dependencies for STEM, and masculinized dependencies for pSTEM (S4 Fig). On one hand, the proportional distribution of the highest level of A&IP (PRIDE D) in the research personnel showed major gender differences (above 0.2) in the Chemistry Institute (STEM) (Figs 3 and 4, S5 and 6 Figs). The same pattern was observed in the Institutes of Chemistry, Cellular Physiology, and Biomedical Research (STEM), where the highest differences in terms of the assignation of Titular C category between male and female names were registered. On the other hand, pSTEM institutes showed in general a more equitable PRIDE D and Titular C distribution (some of them with a higher proportion of female names).

**Fig 3.**
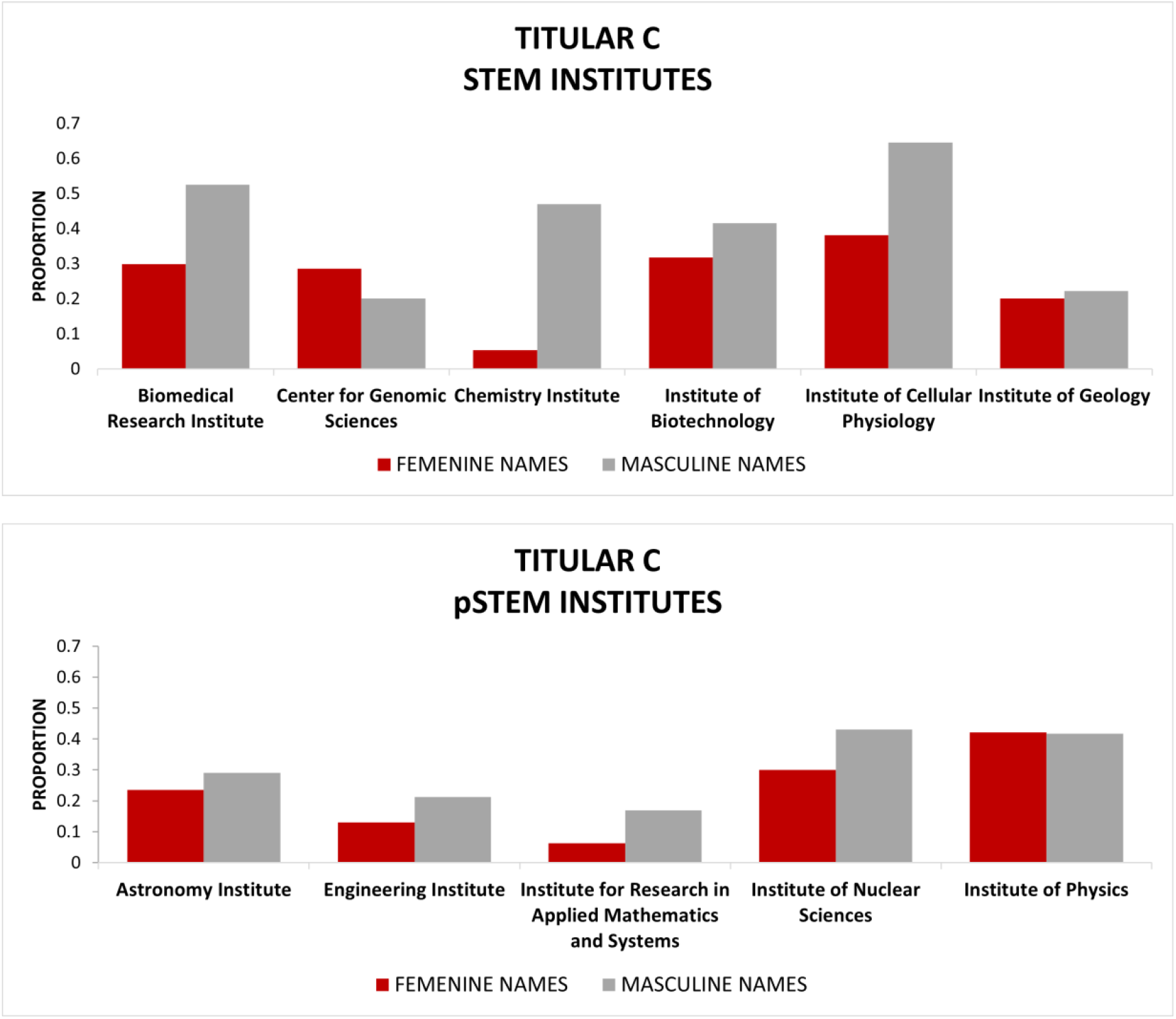
Proportional contribution comparison for the academic position highest level within another STEM and pSTEM UNAM institutes. Feminine and masculine research personnel names are shown. The whole proportional contribution for all levels is shown in S5 Fig. and S6 Fig.

**Fig 4.**
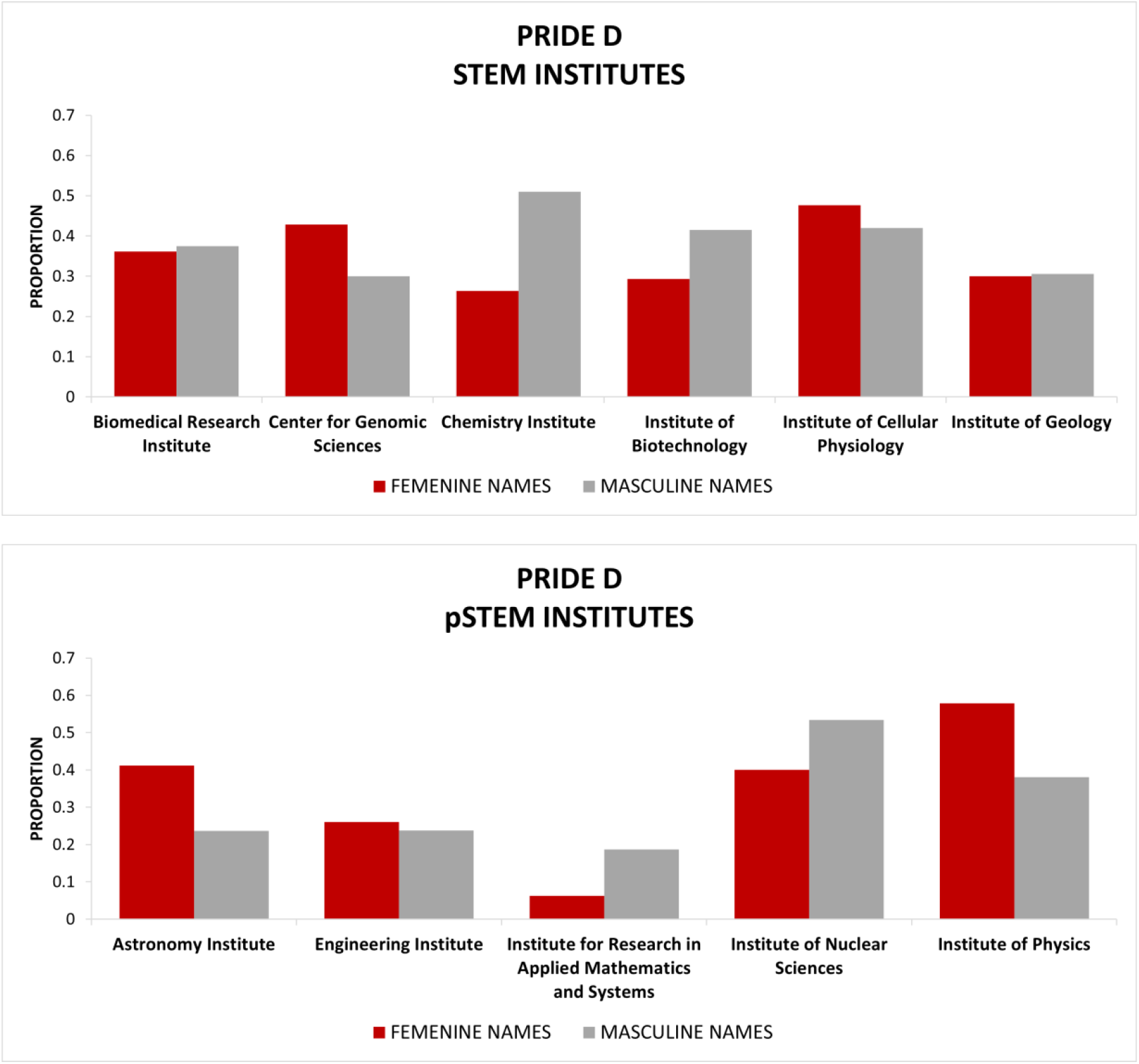
Proportional contribution comparison for the PRIDE incentive program highest level within other STEM and pSTEM UNAM institutes. Same specifications as in Fig 3.

In addition, we observed differences in the average time for promotion in research personnel within the three studied institutes, being higher for the feminine names than for the masculine names (S7 Fig), with some exceptions in the incentive programs (PRIDE and SNI). It is important to emphasize the significant remuneration differences associated with higher levels in both academic and incentive programs (S8 Fig).

## Discussion

### STEM *vs* pSTEM

The analyzed institutions exhibit diverse gender representation, with varying proportions of women and men. The diagnosis of the IM as a pSTEM, matches the unequal distribution of their personnel in terms of gender. STEM disciplines such as biology and ecology are not characterized by horizontal segregation (S1 Table, S4 Fig). At the same time, we detected a significant representation of women in the category of academic technical staff in the IE and IB (S1 Table) and the economic precariousness associated with this category (S2 Table). In this line, the observed masculinization of the academic technical staff within the IM suggests a correlation between the cultural capital of the discipline and traditionally masculine values. This, in turn, underscores the symbolic challenges women face in engaging with this field. That is why, in this case, horizontal segregation is evident, even in the technical-academic staff who have lower incomes).

Regarding academic posts and the phenomenon of horizontal segregation, which is not observed in the IE and IB, we confirmed the accentuation of vertical segregation (Fig 1). In other words, the percentage of women in the highest positions within these two institutes – Titular C and Emeritus - and with the highest incentives - Pride D, SNI III, and SNI Emeritus - is lower than the percentage of men. That is, structural obstacles prevail within these disciplines, whose values are less masculinized and, therefore, have a lower level of cultural capital [11, 15]. We propose that the status in the IE and IB is mainly influenced by the position occupied by each member of the academic staff, and not by the “less-status disciplines” addressed in these institutes. A similar pattern was observed in other UNAM institutes analyzed (Figs 3 and 4), where major differences in masculinization of Titular C and Pride D positions are found in STEM institutes compared with pSTEM institutes.

In the case of the IM, where horizontal segregation is evident (S1 Table), we observed a dilution of vertical segregation. In other words, the percentage of women in the Titular C and with higher incentives - Pride D, SNI III, and SNI Emeritus - (Fig 1 and Fig 2) is similar in almost all cases to the percentage of men. In SNI Emeritus, there are more women. However, we mention that vertical segregation ‘does not disappear’, as reflected in the absence of women Emeritus academic positions and fewer Emeritus women in SNI III (Fig 2), but is greatly diminished in relation to the IE and IB. A dilution of vertical segregation is also observed in other pSTEM UNAM institutes (Figs 3 and 4): Institute of Physics and Institute of Astronomy shows the highest feminization for Pride D position, and pSTEM institutes are in general less masculinized for Titular C position.

This phenomenon, which confirms that horizontal segregation is directly proportional to symbolic barriers and is related to a reduction of vertical segregation, is consistent with a recent meta-analysis that reviewed numerous articles and found that in the United States and Europe, ‘…in tenure-track hiring, our national cohort analyses show no increased likelihood that men proceed to tenure-track jobs relative to women in the very fields in which women are most underrepresented (GEMP), although there is a difference in LPS fields’ (GEMP = geoscience, engineering, economics, mathematics, computer science, and physical science; LPS = life sciences, psychology/behavioral sciences, and social sciences) [4].

At the same time, according to the data from the IE and IB (Figs 1 and 2), we observed vertical segregation. This could be directly proportional to structural barriers, and we found that is associated with more equitable values for horizontal segregation. In these less valued areas - because they are less associated with masculine values and therefore have less cultural capital - the gender hierarchy is reflected in how women are placed in these spaces: these spaces are more precarious because their lower-status posts and incentives result in lower income (S8 Fig). Such female-dominated disciplines have been reported associated with lower success rates [17].

In contrast, when obstacles are symbolic, the hierarchy is reflected in the absence of women and male overrepresentation. In these areas of knowledge with significant cultural capital, the highest posts are less biased for gender reasons: precarization in these cases is shown in the exclusion of women from these types of disciplines. However, we again emphasize that on the one hand, vertical segregation also exists in both STEM and pSTEM institutes (S3, S5 and S6 Figs), but on the other hand, we do observe a lower vertical segregation tendency in pSTEM institutes (Figs 1-4), which was not a blunt pattern. We consider that the context of each institute (internal conditions such as the evaluation criteria for academic staff, which are particular to each one), along with many other structural conditions within each institute are also influencing this tendency.

It is important to mention that we have found, in line with previous studies [8], that women delay their academic promotion in the three institutes (S4 Fig). This reflects a combination of structural and symbolic obstacles. The former obstacle is that women must reconcile their academic-professional life with their family life, a tension not experienced by their male counterparts [18]. This can result in women taking longer to seek category changes; to produce more publications and gain teaching experience, etc. Regarding the symbolic obstacles, women are more insecure and find it harder to trust themselves, a fact that is reinforced by their lower recognition as compared to men [14]. Even in the best case, they must work harder to gain the same recognition as their male counterparts. At this point, which involves subjectivity and qualitative analysis, it is important to investigate how women experience their careers in the three institutes analyzed.

In summary, we consider that this is an innovative study that demonstrates the complex situation of women from science disciplines in a Latin American university. We propose that it is essential to analyze the specificity of the disciplines and their relationship with structural and symbolic obstacles, as well as their relationship with horizontal and vertical segregation phenomena.

We want to emphasize that the solution is not just the inclusion of women in pSTEM; this is necessary, but not sufficient. It is also important to recognize that more women in particular disciplines do not necessarily mean ‘greater equity,’ as demonstrated in IE, IB, and other STEM institutes. At this point, it will be necessary to analyze how women are faring when they are present.

### What is the underlying problem?

This point brings us back to the idea that the problem is not just the underrepresentation of women; therefore, the solution is not simply their inclusion. Instead, by taking the simultaneity of new materialisms as the basis for our analysis [19], we emphasize the overrepresentation of men and their absence in precarized spaces, both physical and symbolic. For example, policies aimed at including women in science have not been matched by policies for including men in communal roles. This discrepancy explains why, between 1995 and 2013, there was an increase in women entering male-dominated occupations, while there was virtually no change in men’s participation in female-dominated fields [15].

In our view, effective inclusion policies should address both structural and symbolic obstacles. Specifically, the goals of these policies should aim to revalorize communal tasks often associated with femininity and promote the inclusion of men in these roles. To achieve this, we must stop perpetuating gender stereotypes that feminize certain tasks and masculinize others through values such as abstraction and rationality.

Furthermore, this study suggests that achieving gender equality is not solely about eliminating horizontal segregation; rather, its reduction seems associated to increased vertical segregation in areas less masculinized. Consequently, the primary aim of inclusion policies should be to ‘de-gender’ the values associated with various disciplines without making them exclusive. De-gendering involves not only ceasing the promotion of stereotypes but also avoiding the devaluation of certain (feminized) occupations while overvaluing others (masculinized). We believe this task is a collective responsibility. If we recognize that caregiving and primary education tasks are predominantly performed by women, we must also acknowledge their role in perpetuating and reproducing gender stereotypes and their associated values. Consistent with our anti-essentialist stance, simply being a woman does not guarantee the end of legitimizing the precariousness of feminized roles or the feminization of precarious positions.

In this study, we advocate for the contextualization and specificity of each discipline. At our university, precarization refers not only to appointments and incentives but also to the distinction between academic technical staff and researchers. One specific policy to address the precarious status of academic technical staff is to revalorize this role in economic, physical, and symbolic terms. We also demonstrate that each institute has unique characteristics influenced by various factors, including the field of knowledge. In this context, stereotypes and forms of precarization also become specific. Although we can make comparisons between institutes, each has distinctive features that deserve qualitative evaluation, and these particularities should be considered in inclusion policies.

## Conclusion

We propose that horizontal segregation in pSTEM could be directly related to the symbolic obstacles that women face and is inversely proportional to the phenomenon of vertical segregation. In contrast, we found a reduction of horizontal segregation in STEM institutes related to Natural Sciences and a higher vertical segregation than pSTEM, which could be linked to structural obstacles.

Fields with high cultural capital create significant symbolic obstacles for women, whereas those with lower cultural capital impose structural challenges. Both types of obstacles --far from being mutually exclusive--coexist, reinforcing a social structure that devalues women.

We must investigate how women may perpetuate stereotypes that keep them away from fields like pSTEM while normalizing their presence in precarized-femenized spaces, leading to their precarization in the academic sphere. This interplay of precarization and feminization perpetuates gender inequality, favoring men through segregation.

Recognizing how the valorization of masculine-associated knowledge contributes to wage disparities becomes indispensable. Addressing this requires examining the interactions across a continuum from communal tasks to academia. As society digitizes, we must question why engineering is valued over caregiving and why mathematical knowledge is prioritized over pedagogical expertise. The prevailing androcentric lens impacts how knowledge is produced and understood, compelling us to redefine values associated with objectivity and neutrality. This involves acknowledging the role of emotions in knowledge production and addressing empirical gaps in discussions around gender differences in emotional processing and abstraction.

## Method

### Data Collection

A systematic search of data related to academic personnel in three scientific research institutes was conducted. The three analyzed STEM institutions included: the Institute of Mathematics (pSTEM), Institute of Biology, and Institute of Ecology, institutions belonging to the National Autonomous University of Mexico (UNAM), characterized by a contrasting gender distribution within the academic personnel: higher numbers of masculine personnel in Mathematics, respecting to Ecology, and Biology. The academic personnel at the UNAM research institutes are grouped into two categories: technical and research personnel. In this paper, we mainly addressed the data corresponding to the research personnel, but we also mentioned some characteristics of technical academic personnel.

A database was constructed based on the following variables: name, gender, age, academic degree, current academic position, academic position history (previous appointments since the incorporation to the institution), current position corresponding to two types of incentive programs, the Academic Personnel Performance Incentive Program (PRIDE, UNAM’s internal program) and the National System of Researchers (SNI, a broad national program), incentives programs history (previous positions in PRIDE and SNI). The gender variable, being protected data, was inferred based on the names of the individuals.

The data collection was performed using different strategies, including: exporting data from public repositories [20], direct requests via the General Transparency Program of the Mexican Government [21], and direct mail petitions to the administration of the analyzed institutes and to General Direction of Academic Personnel (DGAPA) office. DGAPA data 2023 from other UNAM institutes was also requested via the General Transparency Program of the Mexican Government [21]: five pSTEM institutes (Astronomy Institute, Engineering Institute, Institute for Research in Applied Mathematics and Systems, Institute of Nuclear Sciences, Institute of Physics) and six STEM institutes (biological or related areas: Biomedical Research Institute, Center for Genomic Sciences, Institute of Biotechnology, Institute of Cellular Physiology, Institute of Geology). In addition, remuneration data were collected using public repository portals reporting the most recent update (2022 or 2023) related to academic positions [22], and for the incentives programs PRIDE [23] and SNI [24].

### Analysis

A basic descriptive analysis of data was computed by gender (inferred according to the name), as follows: distribution graphs (proportional contribution and age) were obtained considering current data (July 2023) for academic position, and position for incentive programs (PRIDE and SNI) within each institute. To allow a better comparison between institutes --given their differences in feminine and masculine proportion--, we decided to calculate the proportional distribution positions within the academic and incentive programs, according to gender. So, if at IE there are 20 women researchers, and of those 5 are Titular A, 10 Titular B, and 5 Titular C, then they are distributed as follows: 0.25, 0.5 and 0.25 respectively.

To estimate the average time for obtaining an Academic and Incentive Programs (A&IP) promotion, the following was considered: for academic positions, the year and level of the initial hiring category (position in which they began their professional career); for incentive programs, the year and initial level obtained (the first level registered in the database).

Then, the time lapse between one promotion and the next was calculated. Finally, regarding the data related to remuneration, the amount corresponding to research positions, as well as the amount corresponding to incentive programs positions (PRIDE and SNI) for academic personnel were calculated as a percentage concerning the highest A&IP position. The analyses and graphs were done in Excel and with R language version 4.3.

## Acknowledgments

This work was supported by DGAPA, National Autonomous University of Mexico (UNAM) under grant PAPIIT IA400223. This grant PAPIIT IA400223 also funding scholarship of Geraldine Espinosa Lugo and Linda Gisele Zavala Hernández. We thank Nagapriya Wright for his impeccable translation. We appreciate the work of Camila Ramírez Araujo and Adriana Ruiz Gadea for collecting data.

## Supporting information

**Fig S1.**
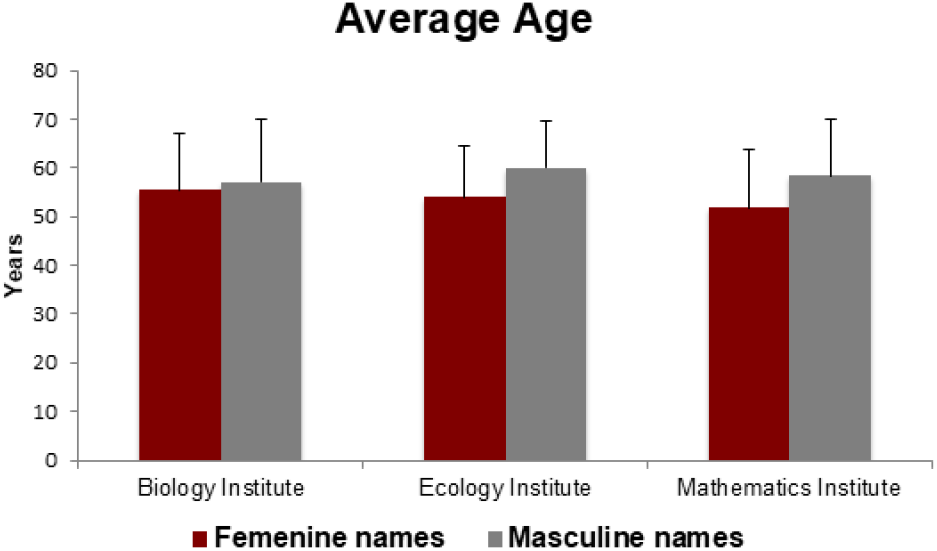
Age average and standard deviation of feminine and masculine research personnel in the three institutes.

**Fig S2.**
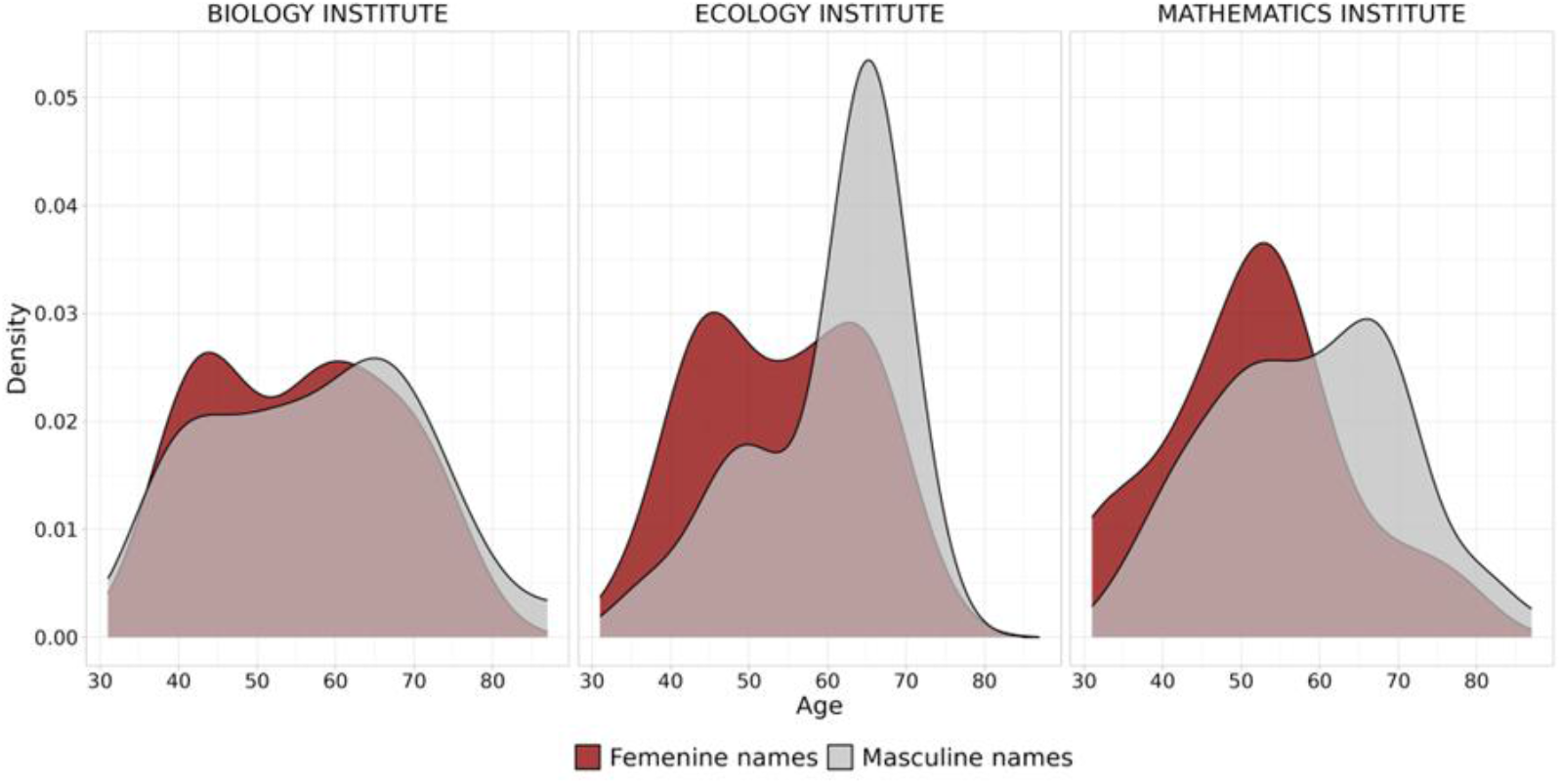
Age normal distribution by gender and institutes (research personnel data).

**Fig S3.**
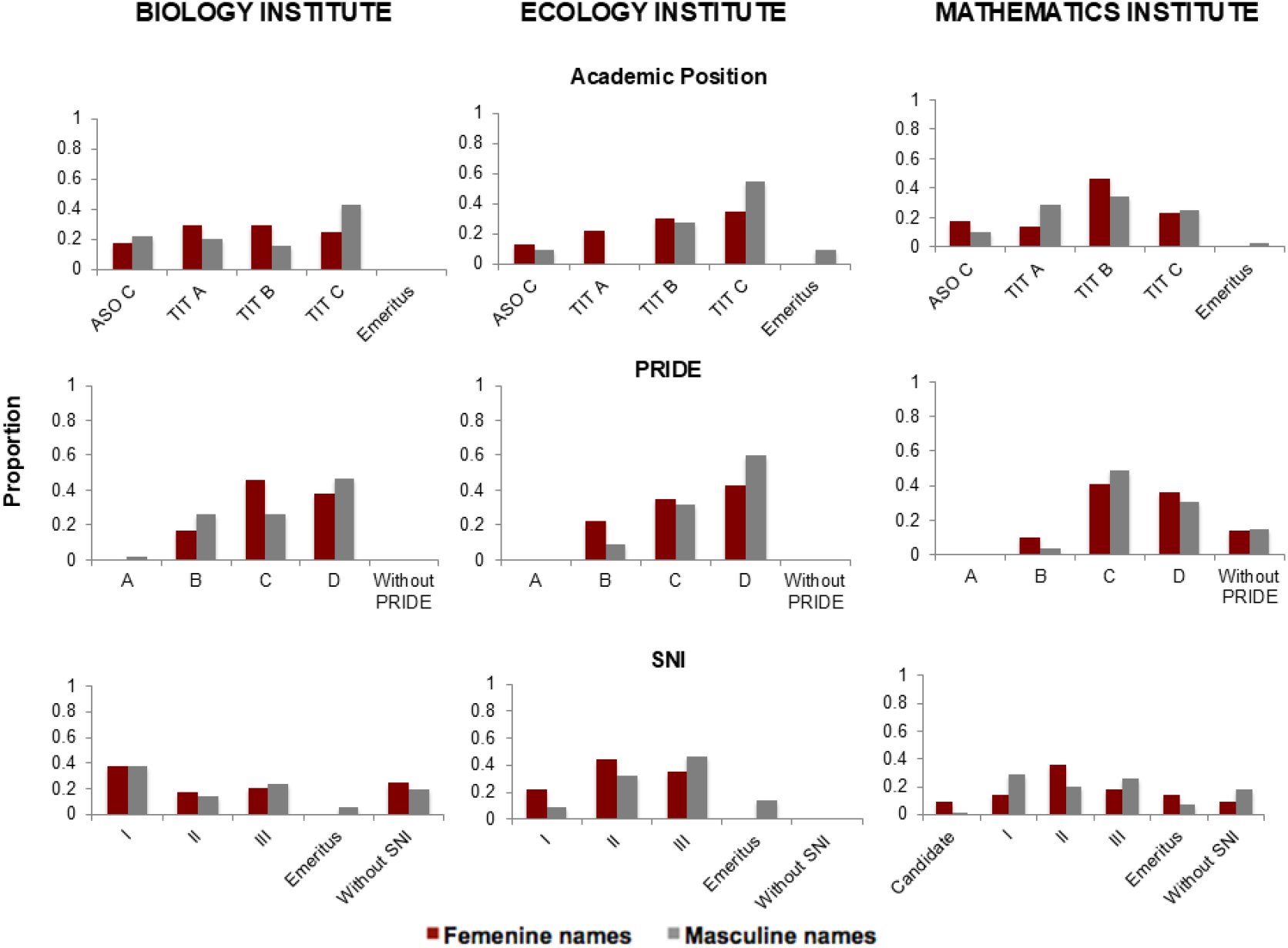
Proportional contribution of feminine and masculine research personnel. From Biology (A, D and G), Ecology (B, E and H) and Mathematics (C, F and I) institutes within academic positions and incentive programs positions.

**Fig S4.**
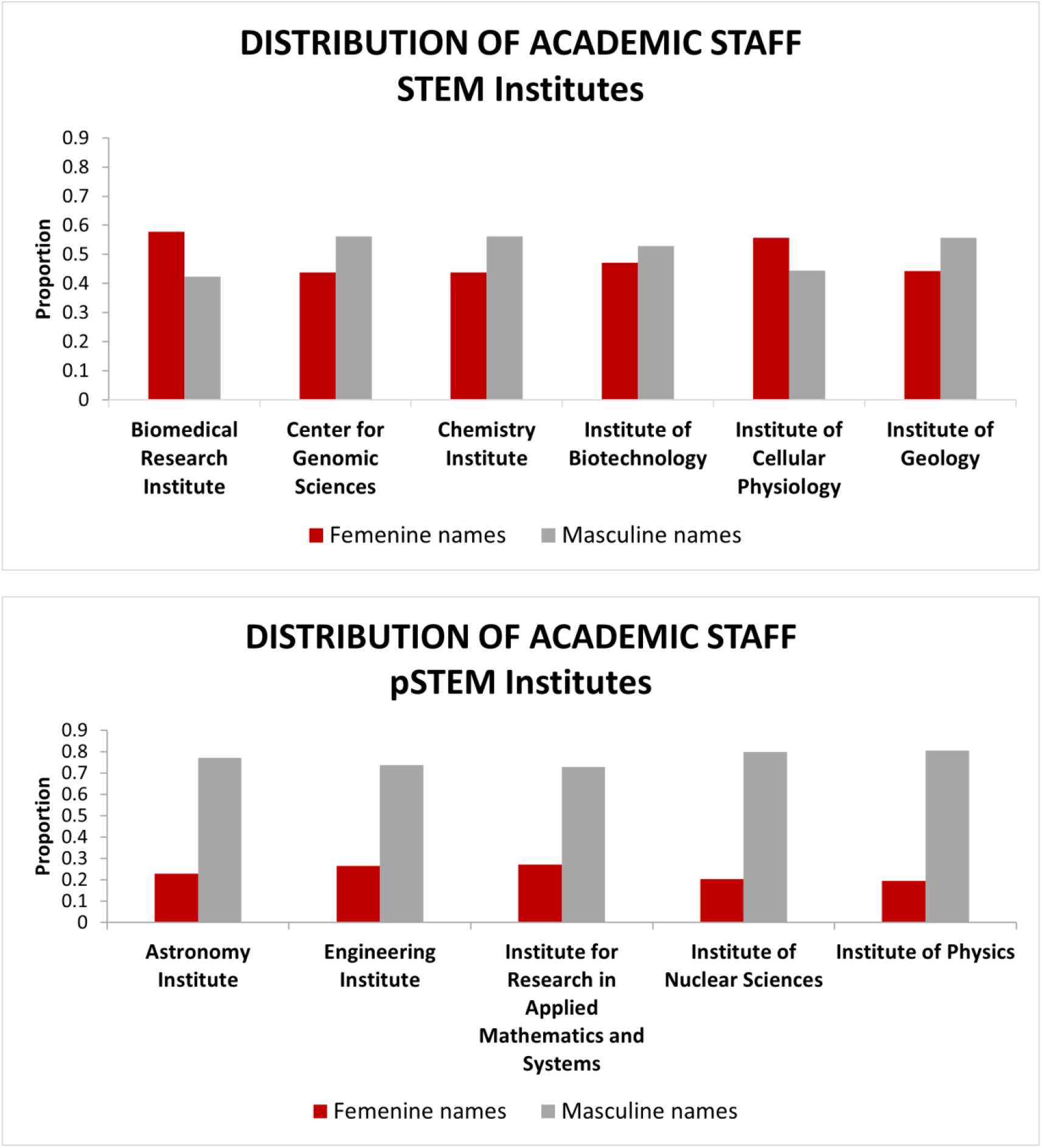
Percentage of masculine and feminine academic personnel names within other STEM and pSTEM UNAM institutes.

**Fig S5.**
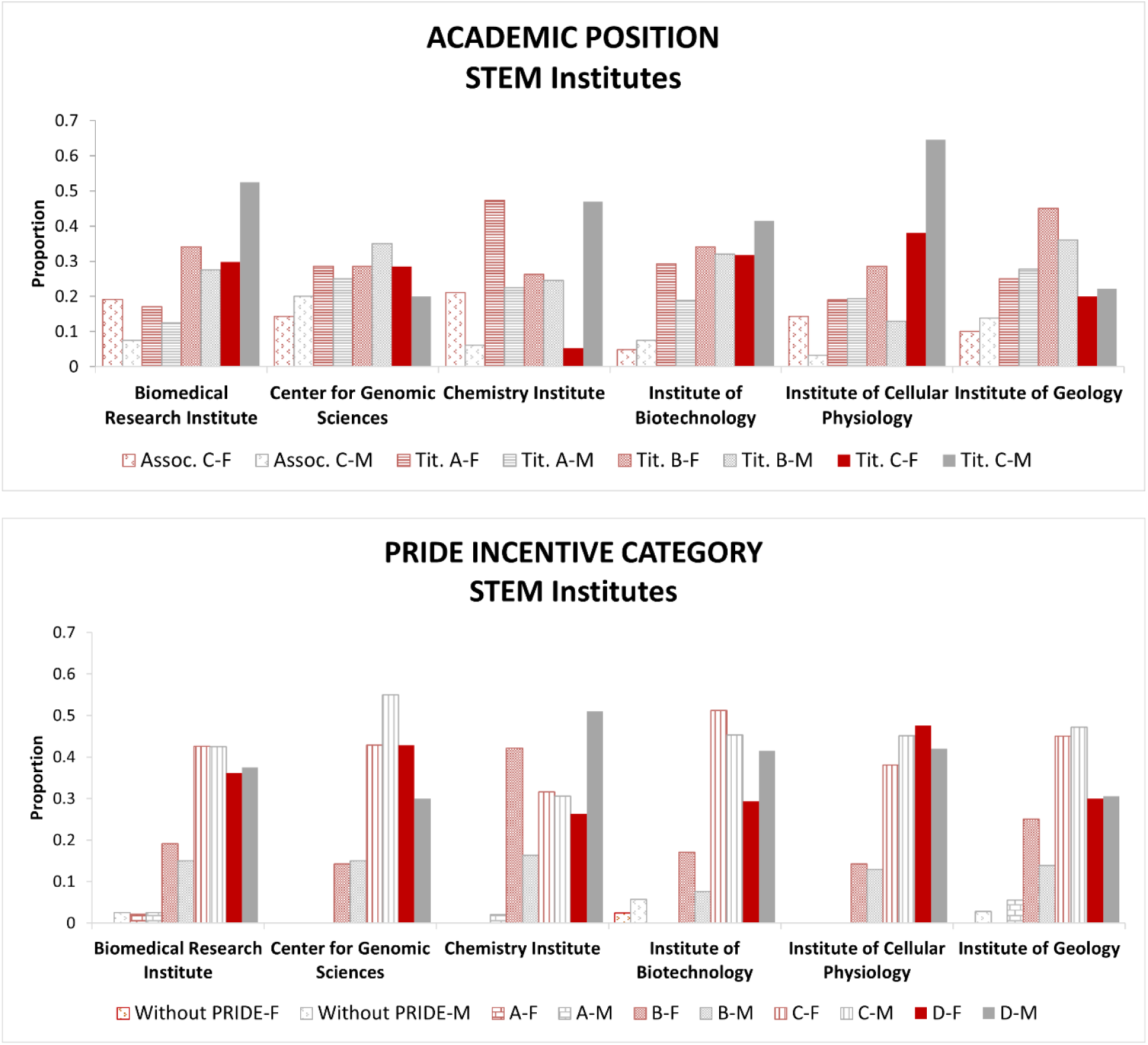
Proportional contribution of feminine and masculine research personnel (additional STEM UNAM institutes) within academic positions and PRIDE incentive program positions. F= Femenine names; M=Masculine names. Assoc. C = Asociado C; Tit A= Titular A; TitB= TitularB; TitC=Titular C; A= Pride A, B = Pride B; C=Pride C; D=Pride D.

**Fig S6.**
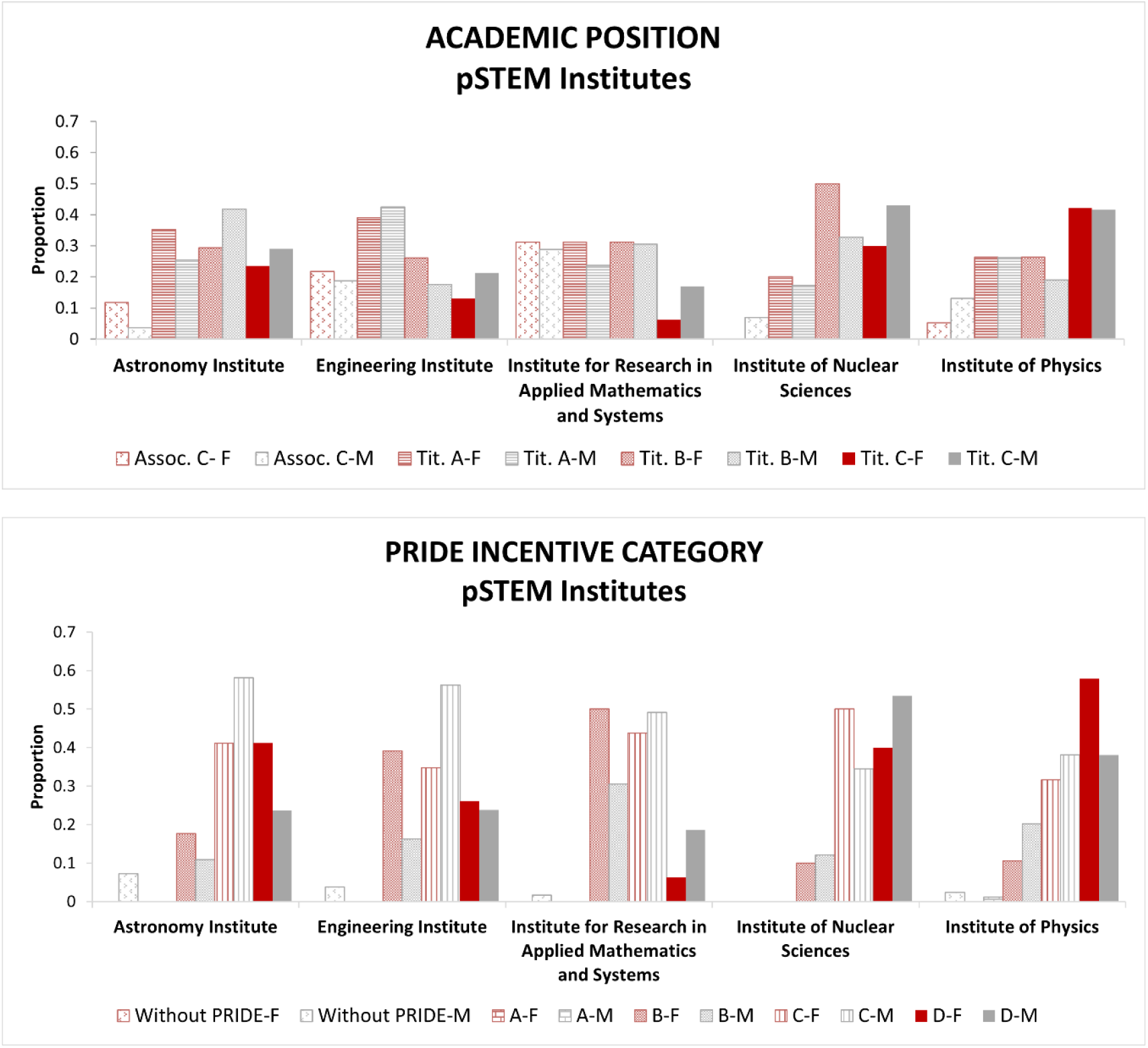
Proportional contribution of feminine and masculine research personnel (additional pSTEM UNAM institutes) within academic positions and PRIDE incentive program positions. F= Femenine names; M=Masculine names. Assoc. C = Asociado C; Tit A= Titular A; TitB= TitularB; TitC=Titular C; A= Pride A, B = Pride B; C=Pride C; D=Pride D.

**Fig S7.**
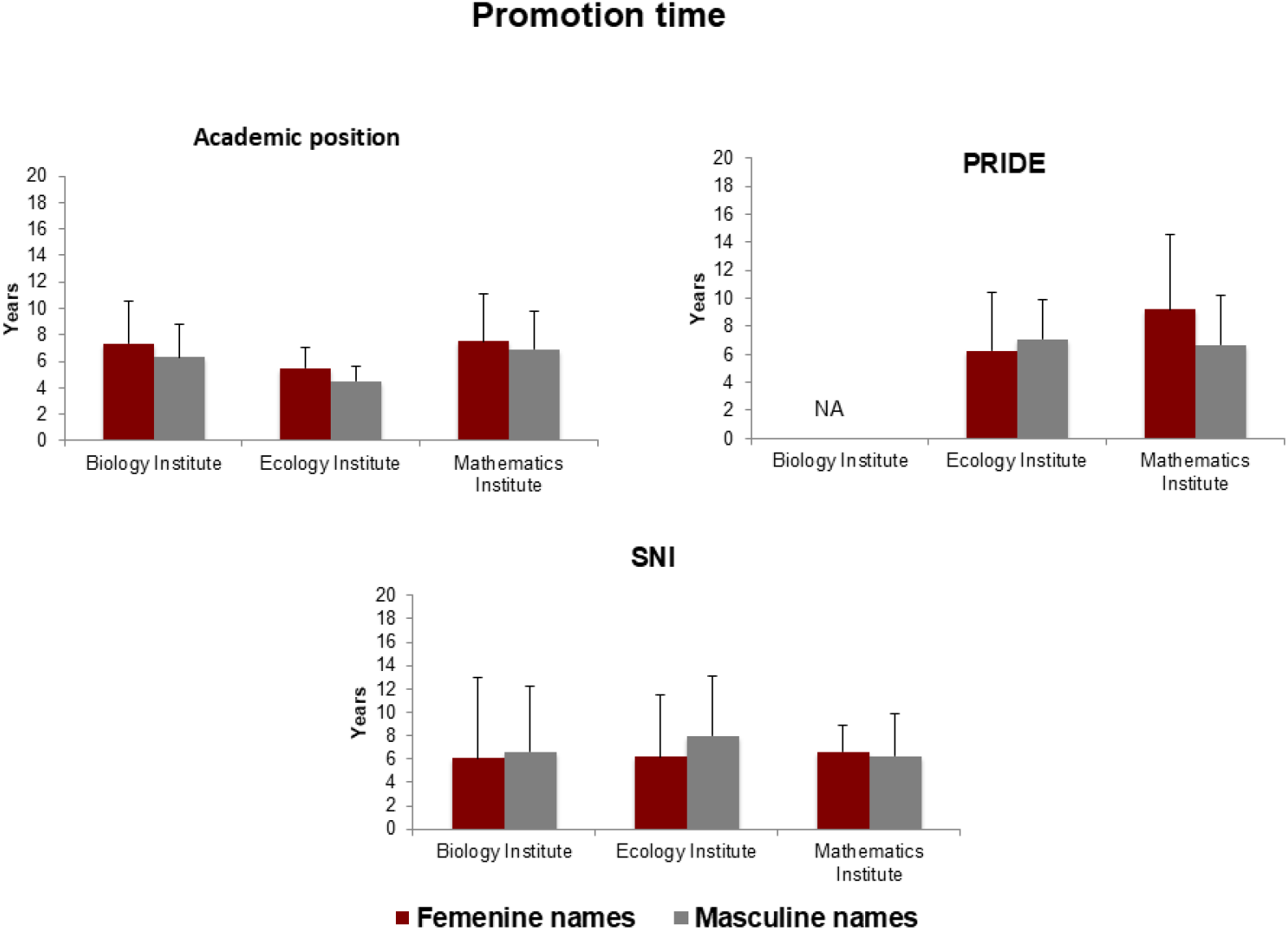
Average time and standard deviation for promotion (years) in feminine and masculine research personnel. Academic (A) and incentive programs (B & C) positions within the 3 institutes analyzed (PRIDE IB data were not available).

**Fig S8.**
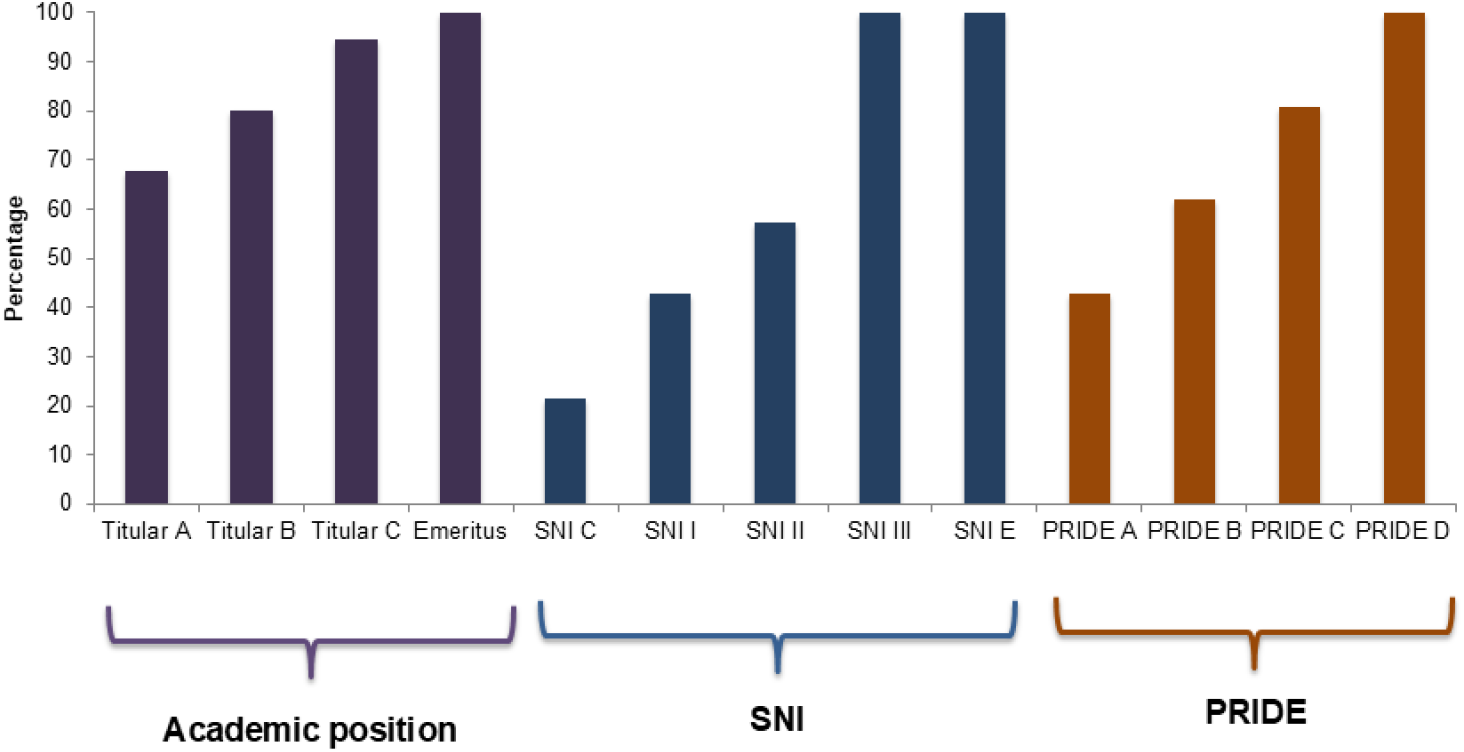
Remuneration differences. Shown in percentage respect to the higher level within academic (research personnel data) and incentive programs positions (IP applies to all academic personnel).

**Table S1.** Feminine and masculine academic personnel counts within the three institutes (IB, Institute of Biology; IE, Institute of Ecology; IM, Institute of Mathematics)

| ACADEMIC PERSONNEL COUNTS |  |  |  |  |  |  |
| --- | --- | --- | --- | --- | --- | --- |
|  | IB |  | IE |  | IM |  |
|  | F | M | F | M | F | M |
| TECHNICAL PERSONNEL | 47 | 39 | 21 | 14 | 6 | 15 |
| RESEARCH PERSONNEL | 24 | 51 | 23 | 22 | 22 | 73 |
| TOTAL | 71 | 90 | 44 | 36 | 28 | 88 |

**Table S2.** 2. Remuneration (mexican pesos) by positions for the academic personnel of UNAM’s institutes corresponding to 2023 year (“Personal UNAM – UNAM Personnel”). The table is ordered from lower to higher levels. NA = Not Applicable (Emerit is a distinction that does not exist for technical personnel). For comparative purposes, we show only the three highest levels for technical personnel (those equivalents to research personnel levels).* *Academic technical staff are responsible for both academic and technical tasks within the university, and there is an implicit devaluation of such roles. The association of ‘technical’ work with productive labour translates into the perception that ‘technical’ activities are of lower status, while academic activities are linked to greater prestige [25]. This devaluation is, of course, associated with economic precarity and can be observed when comparing the perceptions of academic technical staff with those of research staff.

| POSITION | TECHNICAL PERSONNEL | RESEARCH PERSONNEL |
| --- | --- | --- |
| TITULAR A | 19,451.36 | 25,254.4 |
| TITULAR B | 21,882.76 | 29,851.24 |
| TITULAR C | 25,254.4 | 35,287 |
| EMERIT | NA | 37,265.66 |

## References

[1] Dugger WE. Evolution of STEM in the United States. 6Th Bienn Int Conf Technol Educ Res 2010; 1–8.

[2] Merayo N, Ayuso A. Analysis of barriers, supports and gender gap in the choice of STEM studies in secondary education. Int J Technol Des Educ 2023; 33: 1471–1498.

[3] Bøe MV, Henriksen EK, Lyons T, et al. Participation in science and technology: young people’s achievement-related choices in late-modern societies. Stud Sci Educ 2011; 47: 37–72.

[4] Ceci SJ, Kahn S, Williams WM. Exploring gender bias in six key domains of academic science: an adversarial collaboration. Psychol Sci Public Interes 2023; 24: 15–73.

[5] Claybourn C. Majors with the best return on investment. US. News https://www.usnews.com/education/best-colleges/articles/college-majors-with-the-best-return-on-investment, 2022.

[6] McPherson E, Park B. Who chooses a pSTEM academic major? Using social psychology to predict selection and persistence over the freshman year. J Appl Soc Psychol 2021; 51: 474–492.

[7] León LR, Mairesse J, Cowan R. Gender Gaps and Scientific Productivity in MiddleIncome Countries Evidence from Mexico Institutions for Development Sector InterAmerican Development Bank, http://www.iadb.org (2017).

[8] Buquet A. Sesgos de género en las trayectorias académicas universitarias: orden cultural y estructura social en la división sexual del trabajo. Universidad Nacional Autónoma de México (UNAM), 2013.

[9] Harding S. The Science Question in Feminism. Ithaca & London: Cornell University Press, 1986.

[10] Longino H. Subjects, power, and knowledge: description and prescription in feminst philosophies of science. In: Alcoff L, Potter E (eds) Feminist Epístemologies. New York: Routledge, 1993.

[11] Maffia D. Epistemología feminista: La subversión semiótica de las mujeres en la ciencia. Rev Venez Estud la Mujer 2007; 12: 63–98.

[12] Fox Keller E. Reflections on Gender and Science. New Haven: Yale University Press, 1985.

[13] Maffía D. Epistemología feminista: por otra inclusión de lo femenino en la ciencia. In: Blázquez Graf N, Flores J (eds) Ciencia, Tecnología y Género en Iberoamérica. CdMx: México, Centro de Investigaciones Interdisciplinarias en Ciencias y Humanidades, Universidad Nacional Autónoma de México (UNAM), 2005, pp. 623–633.

[14] Maffía D. Conocimiento y emoción. In: Pérez Sedeño E (ed) Monográfico sobre ciencia, tecnología y valores desde una perspectiva de género. Madrid: Arbor, 2005, pp. 516–521.

[15] Croft A, Schmader T, Block K. An underexamined inequality: cultural and psychological barriers to men’s engagement with communal roles. Personal Soc Psychol Rev 2015; 19: 343–370.

[16] Blau FD, Kahn LM. The gender wage gap: Extent, trends, & explanations. J Econ Lit 2017; 55: 789–865.

[17] James A, Buelow F, Gibson L, et al. Female-dominated disciplines have lower evaluated research quality and funding success rates, for men and women. Elife 2024; 13: RP97613.

[18] ciccia lu. La invención de los sexos. Cómo la ciencia puso el binarismo en nuestros cerebros y cómo los feminismos pueden ayudarnos a salir de ahí. Buenos Aires; México; España: Siglo XXI Editores, 2022.

[19] Barad K. Posthumanist performativity: toward an understanding of how matter comes to matter. Signs (Chic) 2003; 28: 801–831.

[20] Tansparencia UNAM UNAM Transparency, http://www.transparencia.unam.mx/sai.html (accessed 1 June 2023).

[21] Portal de Transparencia - Transparency Portal, https://www.plataformadetransparencia.org.mx/ (accessed 1 June 2023).

[22] Personal UNAM - UNAM personnel, http://www.personal.unam.mx/Docs/Contratos/aapaunam-2023_2025.pdf (accessed 1 June 2023).

[23] PRIDE incentive amounts, https://dgapa.unam.mx/images/pride/2023_pride_convocatoria.pdf (accessed 1 June 2023).

[24] SNI incentive amounts, https://dof.gob.mx/nota_detalle.php?codigo=5660859&fecha=10/08/2022#gsc.tab=0 (accessed 1 June 2023).

[25] Moreno H. Ser técnica académica en la UNAM. Colección Estud género y Fem 2022; 27: 11–33.

